# Expression-based machine learning models for predicting plant tissue identity

**DOI:** 10.1101/2023.08.20.554029

**Authors:** Sourabh Palande, Jeremy Arsenault, Patricia Basurto-Lozada, Andrew Bleich, Brianna N. I. Brown, Sophia F. Buysse, Noelle A. Connors, Sikta Das Adhikari, Kara C. Dobson, Francisco Xavier Guerra-Castillo, Maria F. Guerrero-Carrillo, Sophia Harlow, Héctor Herrera-Orozco, Asia T. Hightower, Paulo Izquierdo, MacKenzie Jacobs, Nicholas A. Johnson, Wendy Leuenberger, Alessandro Lopez-Hernandez, Alicia Luckie-Duque, Camila Martínez-Avila, Eddy J. Mendoza-Galindo, David Plancarte, Jenny M. Schuster, Harry Shomer, Sidney C. Sitar, Anne K. Steensma, Joanne Elise Thomson, Damián Villaseñor-Amador, Robin Waterman, Brandon M. Webster, Madison Whyte, Sofía Zorilla-Azcué, Beronda L. Montgomery, Aman Y. Husbands, Arjun Krishnan, Sarah Percival, Elizabeth Munch, Robert VanBuren, Daniel H. Chitwood, Alejandra Rougon-Cardoso

## Abstract

The selection of *Arabidopsis* as a model organism played a pivotal role in advancing genomic science, firmly establishing the cornerstone of today ‘s plant molecular biology. Competing frameworks to select an agricultural- or ecological-based model species, or to decentralize plant science and study a multitude of diverse species, were selected against in favor of building core knowledge in a species that would facilitate genome-enabled research that could assumedly be transferred to other plants. Here, we examine the ability of models based on *Arabidopsis* gene expression data to predict tissue identity in other flowering plant species. Comparing different machine learning algorithms, models trained and tested on *Arabidopsis* data achieved near perfect precision and recall values using the K-Nearest Neighbor method, whereas when tissue identity is predicted across the flowering plants using models trained on *Arabidopsis* data, precision values range from 0.69 to 0.74 and recall from 0.54 to 0.64, depending on the algorithm used. Below-ground tissue is more predictable than other tissue types, and the ability to predict tissue identity is not correlated with phylogenetic distance from *Arabidopsis*. This suggests that gene expression signatures rather than marker genes are more valuable to create models for tissue and cell type prediction in plants. Our data-driven results highlight that, in hindsight, the assertion that knowledge from *Arabidopsis* is translatable to other plants is not always true. Considering the current landscape of abundant sequencing data and computational resources, it may be prudent to reevaluate the scientific emphasis on *Arabidopsis* and to prioritize the exploration of plant diversity.

## INTRODUCTION

Historically, plant biology has focused on inferring genetic, molecular, physiological, and ecological mechanisms. Conventionally, through quantifying phenomena and applying statistics, hypotheses are tested and decisions of most likely scenarios are determined. New technologies and computational approaches have caused a shift from hypothesis- to data-driven research (Mazzocchi, 2015). Moreover, plant biology has embraced the inclusion of machine learning methods in addition to traditional statistical approaches (Ij, 2018). Both a deluge of data and new computational methods have allowed for predictive, rather than inferential, methods. Both statistics and machine learning can be used for inference and prediction, but machine learning methods more often classify and predict on class labels rather than inferring statistical parameters of a population. In plant biology, such predictive approaches underlie the frameworks of phenotyping (Coppens et al., 2017), precision agriculture (Zhang et al., 2002), genomic prediction (Crossa et al., 2014), linking transcriptomic profiles to phenotype (Azodi et al., 2020), and protein structure determination (Jumper et al., 2021). Just as inferential statistics has its limitations, the robustness and ability to extrapolate predictive models are also constrained by the empirical context from which the data originates. Although data-driven research is slowly becoming more theoretical and predictive (Hogeweg, 2011), the creation of universal plant models is hindered by their overwhelming diversity. Not only is the phylogenetic diversity among flowering plants immense (The Angiosperm Phylogeny Group et al., 2016), but plants are exceptionally responsive to their environments (Sultan, 2000) and have evolved symbiotic interactions with and defense mechanisms against innumerable microbes (Mitchell et al., 2006). Furthermore, technical variability in data acquisition makes it difficult to exploit the huge amount of expression data archived in databases. The number of ways we sample molecular profiles from plant tissues and the interaction effects that arise between phylogenetically diverse species with environments, stresses, and biotic interactions is countless and prevents extrapolating results between studies.

Due to the clear advantages of studying a single model species, the early days of the genomics era tended to overlook the importance of prioritizing plant diversity. The candidates considered for the first sequenced genome were either easily transformable (e.g., species within Solanaceae; Knapp et al., 2004) or were already used for genetics (e.g., maize; Strable and Scanlon, 2009), but never was biodiversity considered (Meyerowitz, 2001). Reasons for choosing *Arabidopsis* as the first sequenced plant genome (Arabidopsis Genome Initiative, 2000) include ease of transformation (Clough and Bent, 1998), its small genome (Bennett et al., 2003), and life history traits that allow for genetics through crossing, and short generation times (Meyerowitz, 1987). The justification for initially sequencing the genome of a single model species was that such focus would allow unprecedented molecular discoveries that could be translated into other species and improve our understanding of all plants (Bevan and Walsh, 2005). The strategy to focus on a single model species was successful, and *Arabidopsis* is the most cited plant in the last 20 years, even surpassing key crops and all other plant species (Marks et al., 2023). Our molecular knowledge in plants was purposefully constructed to focus on *Arabidopsis* over crops and plant genetic diversity. However, such a choice has little relevance in a changing climate with dwindling natural resources and vanishing biodiversity that have become the most pressing concerns of our time. The cultural dynamics that influenced the choice of *Arabidopsis* as the first sequenced genome are reflected in subsequently sequenced plant genomes. Plants intrinsic to Indigenous cultures and territories have been sequenced by colonial powers (Marks et al., 2021; Dyer et al., 2022). While sequencing *Arabidopsis* has certainly expanded our knowledge of molecular processes, due to such an intense focus, our understanding in other species remains limited. This leaves us questioning the extent to which the insights from *Arabidopsis* can be extrapolated to the rest of flowering plants.

In the 20 years since the release of the *Arabidopsis* genome sequence (Arabidopsis Genome Initiative, 2000), the number of sequenced plant genomes has dramatically risen (Michael and Jackson, 2013; Li and Harkess, 2018; Marks et al., 2021) leading to a greater understanding of the evolutionary origin and genetic mechanisms underlying numerous traits across the green lineage. Next-generation sequencing, for example, has enabled unprecedented surveys of genome-scale features across species, tissue types, environments, and interactions between plants with abiotic and biotic factors. There are currently over 300,000 public gene expression datasets spanning thousands of diverse plant species (Lim et al., 2022). Cross-species comparisons of gene expression across plants have usually been limited by the number of species analyzed (Proost and Mutwil, 2018) or their sampling breadth. Most studies have generated datasets from scratch (Julca et al., 2021) instead of leveraging public repositories. Databases and datasets curating and making vast amounts of gene expression profiles and their associated metadata have been created. For example, an *Arabidopis* RNA-seq database (ARS) compiles 20,068 publicly available *Arabidopsis* RNA-Seq libraries (Zhang et al., 2020), and the Plant Public RNA□seq Database has ∼45,000 maize, rice, wheat, soybean and cotton samples (Yu et al. 2022). Previously we had curated a dataset of 2,671 publicly available gene expression profiles from 54 flowering plant species across 7 developmental tissue types and nine stresses (Palande et al., 2023). More than 20 years after the release of the *Arabidopsis* genome, not only have we accumulated enough data across plants to ask unprecedented questions but new computational tools are available that permit comparative approaches to analyze such massive amounts of data.

Here, building upon large, curated databases of *Arabidopsis* (Zhang et al., 2020) and flowering plant gene expression profiles (Palande et al., 2023), we examine how predictive *Arabidopsis* is as a model species relative to the rest of the flowering plants and to what degree we can extrapolate our knowledge from model organisms to diverse plant species. Dimension reduction through principal component analysis (PCA) reveals that biotic stress response and tissue type are primary, orthogonal sources of structure in gene expression data from *Arabidopsis*, and while angiosperm data projected onto this space retains some structure, the regions occupied between tissue types become less distinct. We next compare the performance of different machine learning models. The k-nearest neighbor (KNN) method yields precision and recall values of up to 0.99 with models trained and tested on *Arabidopsis* data. Model performance drops significantly, with higher precision than recall values, when data from across flowering plants is tested using models trained on *Arabidopsis* data. Below-ground tissue is more separated from and predictable than other tissue types, and phylogenetic distance from *Arabidopsis* does not appear to influence prediction rates. We end with a discussion of the implications of our results for the current structure of the plant science community, acknowledging that the past focus on *Arabidopsis* as a model organism based on decisions decades ago was effective at that time; however, we now advocate for a shift in approach due to changing circumstances, particularly in light of the pressing issue of biodiversity loss. We argue for a more decentralized and inclusive research framework that better encompasses the diversity of plants and the human cultures that represent them, adapting to current environmental and scientific challenges.

## MATERIALS AND METHODS

### Datasets

We used two curated databases in this analysis. The first contained 28,165 *Arabidopsis* gene expression profiles across 37,334 genes (Zhang et al. 2020). The second contained 2,671 flowering plant expression profiles across 6,327 orthogroups (Palande et al. 2023). Metadata labels for each sample from both of the databases was assigned one of four tissue type labels (above-ground, below-ground, whole plant, or other). The categories are purposefully encompassing and chosen to facilitate accurate assignment across the broad categories of experimental data we analyzed, focusing on above-ground and below-ground tissue identity as one of the simplest cases to test tissue predictability. After removing samples with missing metadata and samples with low unique mapped rate (<75%), the *Arabidopsis* database was left with 19,415 samples. A conserved *Arabidopsis* database was also constructed by keeping only the genes mapped to the orthogroups from the flowering plant database. The conserved *Arabidopsis* database contained the same number of samples, but with much smaller expression profiles across only the 6,327 orthogroups shared with the angiosperm dataset.

### Classification models

Classification is a common machine learning task where, given data points belonging to two or more classes, the goal is to *learn* a function that best differentiates between points from different classes. Then, given a new data point, the function can be used to decide which class the point belongs to. The classifier function can be learned in many different ways, leading to various types of machine learning models. For each classifier model in this study, we employed the following modeling methods:

### Linear support vector classifier (SVC)

In linear classification, each point is viewed as a vector in *k*-dimensional space (Cortes and Vapnik, 1995). The goal is to find *(k-1)*-dimensional hyperplanes that separate the points belonging to different classes. There are many possible choices for hyperplanes that can classify the points. A reasonable choice is to find the ones that maximize the separation between points from different classes. These are known as maximum-margin hyperplanes. Geometrically, the max-margin hyperplanes are defined by the points that lie closest to them; therefore, such points are called support vectors.

### Multi-layer perceptron (MLP)

The SVC model assumes that the classes are linearly separable, which may not be true. MLPs are a class of artificial neural networks (Haykin, 1998) with three or more layers of “perceptrons” with non-linear activation. An MLP consists of an input and an output layer, with one or more hidden layers of neurons. We experimented with one and two hidden-layer MLPs and used rectified linear unit (ReLU) activation in all cases. In ReLU, a neuron ‘s activation is the weighted sum of its inputs, if the sum is non-negative, and zero otherwise. Even with this simple nonlinear activation function, MLPs are able to outperform the linear SVC.

### Random forest (RF)

Random forests (Ho, 1995) perform classification by constructing an ensemble of decision trees. Each decision tree outputs a class label for the given sample and the output of the RF is the class label predicted by the majority of the trees. In a decision tree, each internal node is labeled by an input feature and the leaf nodes are labeled by the class labels. Starting from the root node, the input set is recursively partitioned into children nodes using the input feature associated with the node. The recursion ends when all data points in the node belong to the same class, or some pre-specified termination criteria, such as maximum depth of the tree, are met. Which feature to split the data on at each level is determined using information criteria such as gini impurity or entropy that measure how consistent the subsets are with respect to the class labels after the split.

### Histogram-based gradient boosting (HGB)

Gradient boosting (Mason et al., 1999) is another class of methods that uses a large ensemble of decision trees. In histogram-based boosting, the real-valued input features are first discretized into a few (typically 256) bins using histograms.

This allows the training algorithm to run much more efficiently and construct a much larger ensemble of decision trees to support the classification.

*K-nearest neighbor (KNN) classifier*: In KNN classifiers (Cover and Hart, 1967) class labels are assigned based on a majority vote of the K nearest training points. The distance metric and the number of neighbors are specified by the user. In our experiments, correlation distance between the expression profiles was used to train the KNN classifier.

### Experimental design

To establish the utility of gene expression profiles in predicting tissue type, we trained the supervised machine learning models to classify the *Arabidopsis* data by tissue types (**Table 1**). The database was split into training and test sets (70%-30% split). To ensure comparability, all five models were trained and tested on the same training and test sets. Next, we wanted to examine how predictive *Arabidopsis* is to the rest of the flowering plants (**Table 2**). To test this, we used a set of conserved *Arabidopsis* transcripts with orthogroups across angiosperms, split into training and test sets (70%-30% split) as before. The same five machine learning models were trained on the conserved *Arabidopsis* training set. The performance of these models was first tested on the conserved gene *Arabidopsis* test set to make sure that the models were still able to predict the tissue types with a significantly smaller number of features. We then used the same models to classify the angiosperm data to test how well they extrapolate to species other than *Arabidopsis*. Each machine learning model employed in our experiments requires additional hyperparameters that need to be tuned to optimize model performance. We used the Bayesian optimization procedure implemented in the hyperopt package in Python (Bergstra et al., 2013). To gain insights into the functional annotation and enrichment of our gene list, we performed a Gene Ontology term analysis using the DAVID Functional Annotation Clustering tool (version 2021) from the web interface http://david.ncifcrf.gov (Huang et al., 2009). We filtered the 200 genes with the most positive and negative PC1 loading values. The annotation was performed using TAIR IDs and selecting Gene Ontology terms from levels 3 and 4 of Molecular Function and Biological Process categories. All data and code to reproduce the results in this manuscript are available at https://github.com/PlantsAndPython/arabidopsis-gene-expression.

**Table 1:**
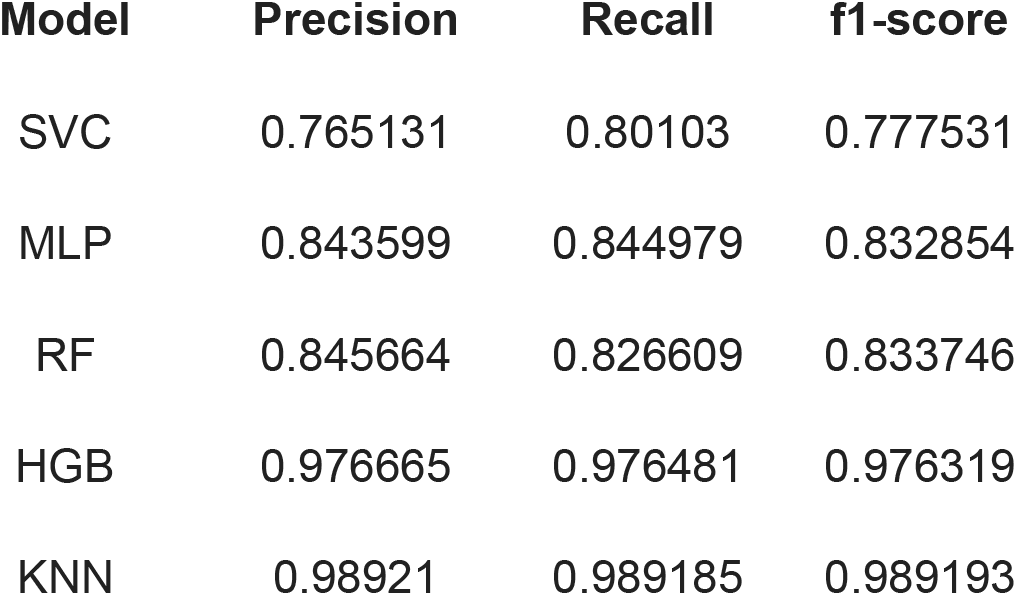
Classification performance of models trained on the full *Arabidopsis* dataset.

**Table 2:** Classification performance of models trained on the conserved *Arabidopsis* dataset and tested on conserved *Arabidopsis* or Angiosperm datasets.

| Model | Test Set | Precision | Recall | f1-score |
| --- | --- | --- | --- | --- |
| SVC | Arabidopsis | 0.740855 | 0.778026 | 0.754276 |
|  | Angiosperm | 0.695691 | 0.576189 | 0.591683 |
| MLP | Arabidopsis | 0.822682 | 0.828155 | 0.824351 |
|  | Angiosperm | 0.734603 | 0.547361 | 0.611767 |
| RF | Arabidopsis | 0.862941 | 0.864721 | 0.861927 |
|  | Angiosperm | 0.747272 | 0.569075 | 0.622122 |
| HGB | Arabidopsis | 0.971034 | 0.970987 | 0.970574 |
|  | Angiosperm | 0.741902 | 0.567952 | 0.640741 |
| KNN | Arabidopsis | 0.987804 | 0.987811 | 0.987803 |
|  | Angiosperm | 0.733478 | 0.643205 | 0.663313 |

## RESULTS

### Dimension reduction and alignment between Arabidopsis and angiosperm gene expression datasets

A principal component analysis (PCA) performed on the full dataset of 19,415 *Arabidopsis* RNAseq samples shows a clear separation by tissue type (**Fig. 1a**). For simplicity, we categorized samples into bins of above-ground, below-ground, whole plant, and other. The above-ground, below-ground, and other tissue types are well-separated from each other, but the below-ground tissue has the least overlap with other tissues. The whole plant tissue type, composed of different combinations of the other tissues, is not well separated, as we would expect. The separation of tissues occurs along a gradient defined by PC2, demonstrating that tissue type is not the primary source of variance in the data. Rather, a small proportion of samples are strewn across PC1 in an additive, orthogonal manner, preserving the separation of tissue types defined by PC2. To investigate the underlying cause responsible for the primary source of variation in the data, we performed GO enrichment on genes with the most extreme PC1 loading values that are most responsible for defining PC1. In the full *Arabidopsis* dataset (**Fig. 1a**), high PC1 values, which include a small number of samples that contribute to a disproportionate amount of variance in the data, are defined by high expression of genes associated with response to biotic stress and oxidative damage GO terms (**Table S1**). Low PC1 values, which include a majority of samples across tissues and which we assume arise from plants grown under regular conditions associated with the less stress, are defined by high expression of genes with GO terms associated with biosynthesis, biogenesis, and cell growth. Remarkably, in the full *Arabidopsis* dataset, negative PC1 loading values are enriched for glucosinolate biosynthetic and other metabolic processes (FDR <0.05).

**Figure 1:**
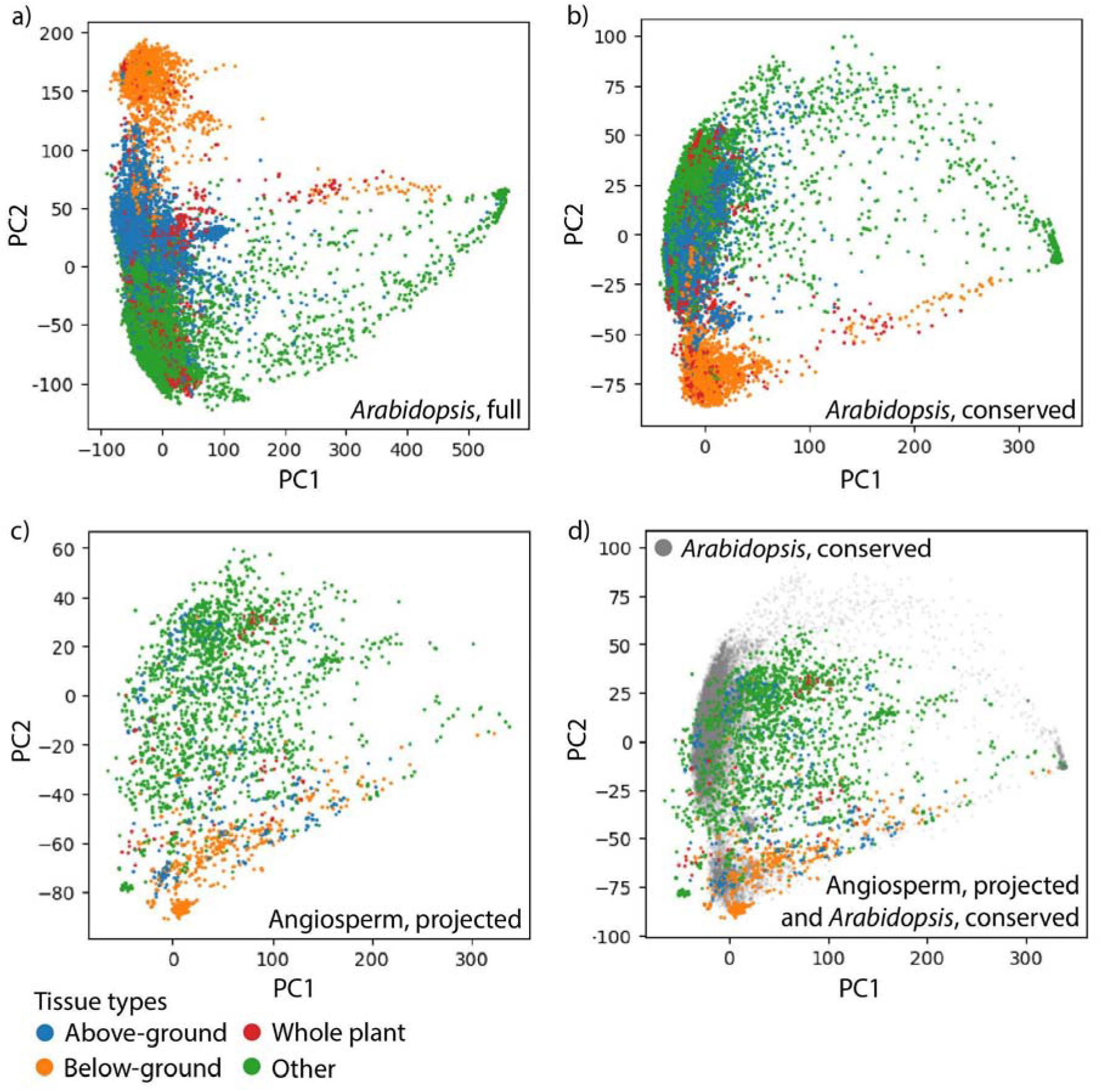
Principal Component Analysis (PCA) of gene expression profiles. PCAs with gene expression profiles colored by above-ground (blue), below-ground (orange), whole plant (red), and other (green) tissue types for **a)** the full *Arabidopsis* dataset, **b)** the conserved *Arabidopsis* data set, **c)** the angiosperm dataset projected onto the conserved *Arabidopsis* PCA from b), and **d)** the same as c), but with conserved *Arabidopsis* gene expression profiles in the background (transparent gray).

From these large-scale datasets, we developed a predictive model to test if tissue type could be inferred from expression patterns alone and if this *Arabidopsis*-trained model could be transferred to other flowering plants. We previously created a set of 6,328 low copy orthogroups that are deeply conserved across flowering plants (Palande et al., 2023) and used a set of 6,327 *Arabidopsis* genes corresponding to these orthogroups for all downstream analyses. A PCA performed on this subset of 6,327 conserved flowering plant genes shows mostly the same structure as the analysis with all *Arabidopsis* genes included (**Fig. 1b**). However, while the below-ground tissue type remains distinct from the rest of the data, the above-ground tissue type overlaps more with whole plant and other tissue types. Note that the sign of principal components is arbitrary, which explains the “flip” of PC2 values relative to the full set of *Arabidopsis* genes. An analysis of the enriched GO terms for PC1 loading values from the conserved gene PCA reveals that high PC1 values are associated with biotic responses, but also with anther- and pollen-related GO terms (**Table S1**). Low PC1 values are associated overwhelmingly with photosynthesis. Because the two datasets have corresponding orthogroup features, we are able to project the angiosperm dataset onto the PCA defined by the conserved gene *Arabidopsis* dataset (**Fig. 1c-d**). While the overall structure defining the distributions of tissue types is maintained in the projected angiosperm data, there is substantial overlap between above-ground and below-ground tissue types. We conclude that indeed there is conservation of tissue-specific expression between *Arabidopsis* and the rest of the flowering plants, but that as expected, the alignment of the underlying structures of gene expression patterns defining tissue type identity are not identical.

### Predictive modeling of plant tissue from gene expression

We used supervised learning classifiers to test if gene expression profiles could predict tissue type in *Arabidopsis* and if these *Arabidopsis* trained models could be applied more broadly to flowering plants. We first split the *Arabidopsis* data into testing and training sets with samples split into four classes of above-ground, below-ground, whole-plant, or other as described above. Models trained on *Arabidopsis* expression data and used to predict tissue type in *Arabidopsis*, whether the full or conserved gene datasets, achieved high precision and recall scores. The highest f1-scores (the harmonic mean of precision and recall) for the full and conserved datasets were achieved using a K-Nearest Neighbors algorithm (KNN) (0.99 and 0.99, respectively; **Tables 1 and 2**) and the lowest using Linear Support Vector Classification (SVC) (0.78 and 0.75). Histogram-Based Gradient Boosting (HGB) also achieved high f1-scores (0.98 and 0.97) while the results for Random Forest (RF) (0.83 and 0.86) and Multilayer Perceptron (MLP) (0.83 and 0.82) were intermediate. When used to predict *Arabidopsis* data, the precision and recall values for each model were similar to each other, indicating similar positive prediction value (precision, true positives divided by true positives and false positives) and sensitivity (recall, true positives divided by true positives and false negatives). The relative prediction rates of different tissue types to each other were equivalent for the full *Arabidopsis* dataset (**Fig. 2a)**.

**Figure 2:**
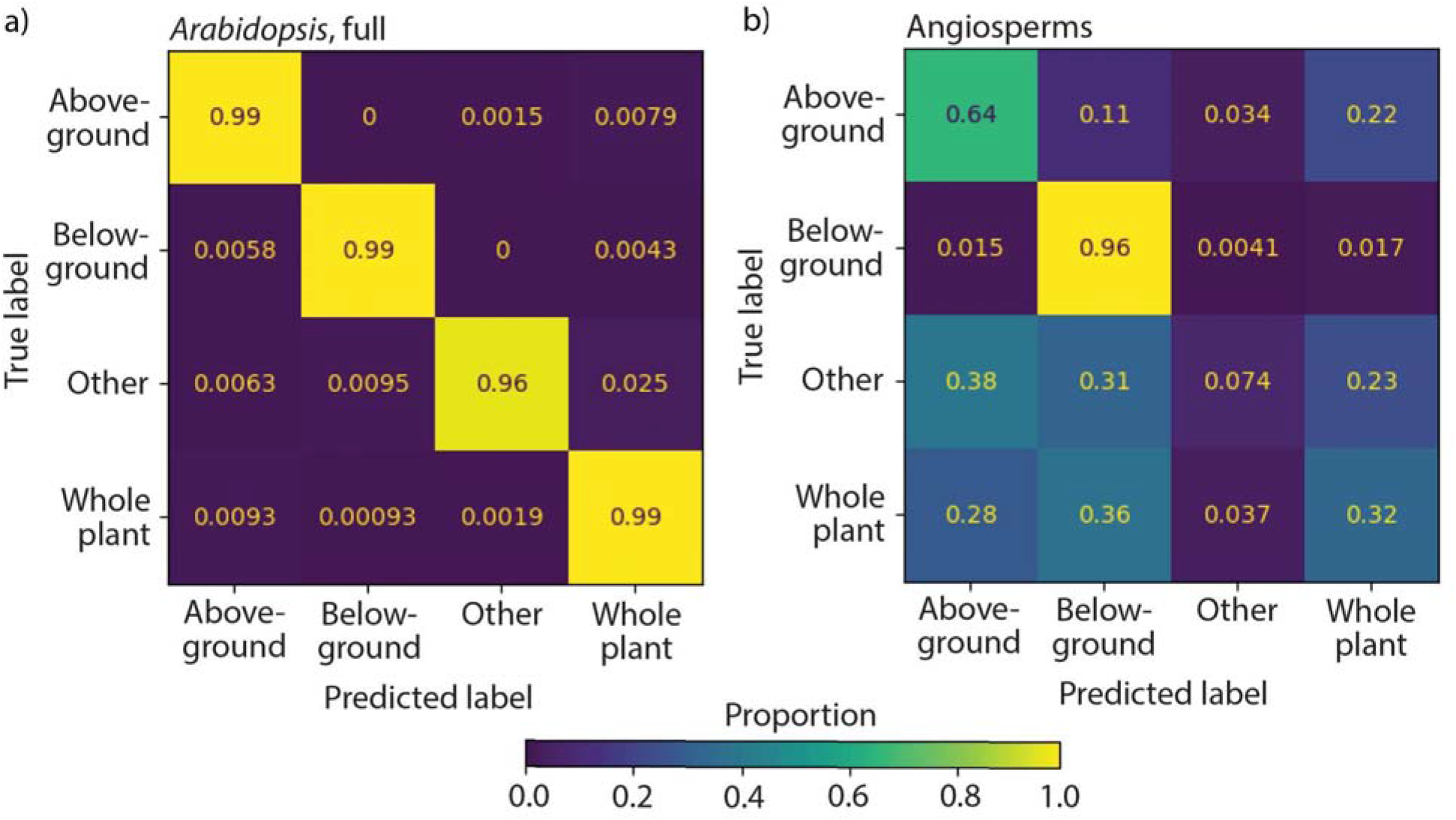
Confusion matrices using the KNN-classifier. Confusion matrices showing true label identity (vertical axis) and the proportion of samples assigned to predicted label identities (horizontal axis) for **a)** the full *Arabidopsis* dataset and **b)** the angiosperm dataset. Proportion indicated by viridis color scale.

The projection of gene expression patterns from across flowering plants onto a PCA using a conserved set of genes from *Arabidopsis* shows considerable variability (**Fig. 1c-d**). Using models trained on *Arabidopsis* data and tested on flowering plants, prediction rates are more similar to each other using different algorithms than *Arabidopsis* alone but perform much worse, and with higher precision than recall rates (**Table 2**). For KNN, HGB, RF, MLP, and SVC methods, precision values were 0.73, 0.74, 0.75, 0.73, and 0.70, respectively, whereas the rates of recall were 0.64, 0.57, 0.57, 0.55, and 0.58. Although these rates are moderately high, they must be interpreted in the context of using only four tissue type labels. The relatively higher precision rates compared to recall indicate that when a sample is retrieved, there is a higher rate of the models calling a true positive (positive prediction value) compared to the fraction of relevant samples retrieved (sensitivity). The prediction rates across tissue types were not evenly distributed (**Fig. 2b**). Below-ground tissue was accurately classified, at a rate of 0.96, while above-ground tissue was only correctly predicted at a rate of 0.64. Other and whole plant tissue types were classified poorly (0.074 and 0.32, respectively), and almost no samples were predicted as other tissue type, including other samples themselves. Although the prediction accuracy varies considerably across plant families (**Fig. 3**), from around 0.4 to 0.8, we could not identify any phylogenetic signal or find any support that prediction of tissue identity is inversely correlated with distance of a plant family from *Arabidopsis* in the Brassicaceae.

**Figure 3:**
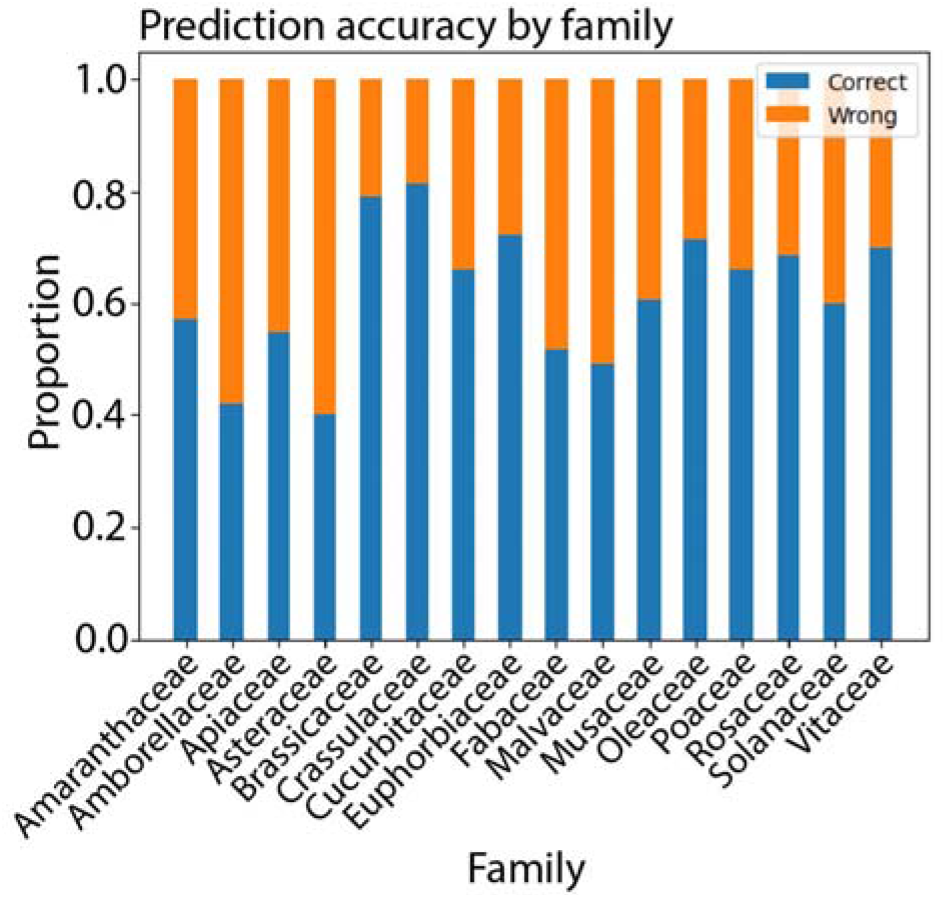
Prediction accuracy by plant family. Using KNN-classifier on the angiosperm dataset, the proportion of samples correctly (blue) and wrongly (orange) predicted from *Arabidopsis* data is shown as a stacked bar plot.

## DISCUSSION

### Arabidopsis-only models are highly accurate

Although we focus on tissue identity in this study, we note that the strongest source of variance (PC1) in publicly available *Arabidopsis* gene expression profiles is a signature associated with biotic defense (**Table S1**) and that it acts in an additive, orthogonal manner with respect to tissue type which is the next strongest source of variance (PC2). Not only are higher prediction rates expected for the *Arabidopsis-*only models because the same dataset is being used for training and testing, but because of the data structure itself that separates the main factors we are testing— above and below ground tissues—as visualized in a PCA (**Fig. 1a-b**). From this perspective, it is perhaps not surprising that KNN is the best performing algorithm, based on the overall distance-based proximity of gene expression profiles for each label to each other (**Table 1**). The other methods, based on decision trees or neural networks, by focusing on individual gene expression values as parameters, fail to account for overall distance. The focus on individual gene expression values instead of the overall signature or profile is reminiscent of the molecular biology concept of “biomarkers” to indicate the tissue or stress from which a sample arises. The outperformance of KNN over other algorithms we tested may suggest that gene expression signatures (rather than focusing on individual gene expression values) are more valuable to create models for tissue and cell type prediction.

### Arabidopsis gene expression as a model for other flowering plants may not be the most suitable approach

Lower prediction rates are expected when testing a model on different data than its training set (**Table 2**). However, the lower precision and recall scores when a model trained on *Arabidopsis* is tested on gene expression samples across the flowering plants undermines the foundational argument for using model species: that data from *Arabidopsis* would be predictive for plants in general. This is not to say that there is not substantial conservation of tissue-specific gene expression patterns. Our own work (Palande et al., 2023) and that of others (Julca et al., 2021) strongly supports conserved tissue-specific gene expression patterns across flowering plants, as is true of animals as well (Fukushima and Pollock, 2020). Rather, the ability to leverage and predict tissue identity from conserved gene expression profiles is diminished when building a model from a single, arbitrary species.

Details of the performance of our model hint at underlying biological considerations when using model species data. Not all tissue types are equally predictable, and the prediction of below-ground tissue outperforms other tissue types (**Fig. 2**). We hypothesized that the ability to predict tissue identity from *Arabidopsis* may be inversely correlated with phylogenetic distance of a sample from Brassicaceae, but we found no evidence to support this idea (**Fig. 3**). Additionally, the precision values for predicting tissue type of flowering plant data from *Arabidopsis* are much higher than recall values (**Table 2**). This may indicate that models are relatively better at calling samples with conserved tissue-specificity with *Arabidopsis* (a true positive) over those without (a false negative). These results may also be a product of our classification scheme. For example, above and whole plant tissues are often more similar to each other than below ground tissue because they are missing roots, and might more easily be misclassified with each other. The other category is composed of diverse tissues which may not have clear predictive features. These factors should be considered when evaluating the classification results (**Fig. 2**).

Our results potentially arise not only from genes with evolutionary differences in tissue-specific expression compared to *Arabidopsis*, but ones that may indeed have conserved expression but differ in the ways we have culturally constructed our developmental descriptions of plant species. Such a circumstance might arise when the cell type-specific expression of a gene is truly conserved, but that evolved differences in functional morphology between species lead us to apply different tissue descriptors (for example, between an herbaceous annual and a woody perennial, or a CAM succulent compared to a weedy C3 plant). The misalignment of tissue labels extends to more quantitative descriptors and to the molecular level, including Gene Ontology (GO) and Kyoto Encyclopedia of Genes and Genomes (KEGG) terms that ultimately become biased to plants with sequenced genomes (Provart et al., 2016). For example, in our analysis of genes corresponding to the most positive and most negative PC1 loading values, there was a noticeable enrichment of genes associated with the glucosinolate biosynthetic and metabolic pathways in *Arabidopsis* samples (**Table S1**). However, this enrichment was absent in broader angiosperm samples, as these compounds are found almost exclusively in Brassicaceae. Glucosinolates are a diverse group of secondary metabolites that play a critical role in plant defense against herbivores and pathogens. Beyond their defensive role, they seem to be involved in growth, development, microbiota interactions, and phosphate nutrition (Kopriva, 2021). Focusing on a single organism, or small group of model species to predict attributes of all plants is flawed from both biological (arising from evolutionary novelty) as well as philosophical (due to semantic, ontological, and cultural differences in how we socially construct plants) perspectives.

### Moving forward and embracing plant and cultural diversity

*Arabidopsis* was selected as a model species unilaterally, over raised objections, decades ago arising from mostly genetic and molecular biology considerations (Meyerowitz, 1987; Clough and Bent, 1998; Arabidopsis Genome Initiative, 2000; Bennett et al., 2003; Bevan and Walsh, 2005). Arguments in favor of plant diversity or selecting agricultural or ecological models were ignored. These past decisions have led to continued focus on *Arabidopsis* and there is continuing advocacy for a plant model species and to fund *Arabidopsis* research at the expense of plant diversity to this current day (Provart et al., 2016; Parry et al., 2020). Since then, data science and computational approaches have begun to grow. Retrospectively, after which decades of sequencing data across flowering plants has allowed us to objectively ask if the focus on a single, arbitrary plant allows us to predict the biology of other flowering plants better than if we had studied all plants equally from the start, the answer is no (**Table 2**). Using a data science approach and building machine learning models on *Arabidopsis* gene expression data to predict the tissue identity of gene expression samples from across flowering plants as we have done here, does not preclude the consideration of other, more important qualitative arguments against the model species concept that continue to limit the potential of the plant science community. Beyond just *Arabidopsis*, there is still a focus on agriculturally important species at the expense of all plants (Marks et al., 2023). More insidiously, the social construct of plants and their diversity arises from colonialism, evidenced not only by the gaze of the Global North and the plants we have chosen to research and document and how we do so, but in ways that can be quantified related to the specific discussion of *Arabidopsis* here, specifically which plant genomes have been sequenced and by whom (Marks et al., 2021), usually through extinguishing and stealing the cultural knowledge of Indigenous people (Dwer et al., 2022).

Useful discoveries and insights have arisen from *Arabidopsis* (Arabidopsis Genome Initiative, 2000). Rather than advocating for continued focus and funding for a single model species (Provart et al., 2016; Parry et al., 2020), it is long past due that we address the historical inequities that have led to our current construction of the plant sciences and that we avoid a biased focus and embrace the biological and cultural diversity of the plant world.

## Data availability

The code, metadata, and raw datasets from this project are available on a dedicated GitHub page: https://github.com/PlantsAndPython/arabidopsis-gene-expression

## Acknowledgements

This work was funded primarily by an NSF-NRT training grant (NSF 1828149) which established the Integrated training Model in Plant And Compu-Tational Sciences (IMPACTS) program at Michigan State University. This grant funded fellows within this program as well as the project based curriculum for the Plants and Python Course that formed the backbone of this manuscript. This work is also supported by NSF Plant Genome Research Program awards IOS-2310355, IOS-2310356, and IOS-2310357, and NSF Plant, Fungal and Microbial Developmental Mechanisms award IOS-2039489. This project was supported by the USDA National Institute of Food and Agriculture, and by Michigan State University AgBioResearch.

